# Impaired trigeminal control of ingestive behavior in the Prrxl1^- / -^ mouse is associated with a lemniscal-biased orosensory deafferentation

**DOI:** 10.1101/2021.10.07.463562

**Authors:** Admir Resulaj, Jeannette Wu, Mitra J. Z. Hartmann, Paul Feinstein, H. Phillip Zeigler

## Abstract

Although peripheral deafferentation studies have demonstrated a critical role for trigeminal afference in modulating the orosensorimotor control of eating and drinking, the central trigeminal pathways mediating that control, as well as the timescale of control, remain to be elucidated. In rodents, three ascending somatosensory pathways process and relay orofacial mechanosensory input: the lemniscal, paralemniscal, and extralemniscal. Two of these pathways (the lemniscal and extralemniscal) exhibit highly structured topographic representations of the orofacial sensory surface, as exemplified by the one-to-one somatotopic mapping between vibrissae on the animals’ face and barrelettes in brainstem, barreloids in thalamus, and barrels in cortex. Here we use the Prrxl1 knockout mouse model to investigate ingestive behavior deficits associated with disruption of the lemniscal pathway. The Prrxl1 deletion disrupts somatotopic patterning and axonal projections throughout the lemniscal pathway but spares patterning in the extralemniscal nucleus. Our data reveal an imprecise and inefficient ingestive phenotype with deficits that span timescales from milliseconds to months, tightly linking trigeminal input with ingestion, from moment-to-moment consummatory to long term appetitive control. We suggest that ordered assembly of trigeminal sensory information along the lemniscal pathway is critical for the rapid and precise modulation of motor circuits driving eating and drinking action sequences.

## 1. Introduction

In rodents, sensory information from the orofacial region is critical for the moment-to-moment control of two effector systems centrally involved in ingestive behavior: the whiskers and the mouth. Active sensing by the whiskers is important for appetitive behaviors, including the localization and identification of food and water sources. Inputs from the perioral, oral and intraoral regions modulate consummatory behaviors, including the grasping, manipulation, and licking movements involved in eating and drinking.

Both sets of orofacial inputs are conveyed to the brain by the trigeminal (V) nerve, whose cell bodies reside in the V ganglion (Vg) and branch to innervate the entire brainstem trigeminal complex, including the principal and spinal trigeminal nuclei (PrV and SpV, respectively). PrV originates the lemniscal pathway, which relays through the dorsomedial portion of the ventral posteromedial thalamus (VPMdl) to terminate in layer IV of primary somatosensory cortex (S1). SpV originates two pathways: the paralemniscal, which starts in SpVir, continues to the posteromedial complex of the thalamus (PoM), and terminates in secondary somatosensory cortex (S2), vibrissal motor cortex, and layer Va of S1; and the extralemniscal, which starts in SpVic, passes through the ventrolateral regions of (VPMvl) and continues to S2 and layer Vb of S1 (1-3).

Cytochrome oxidase (CO) staining reveals that several of these regions (PrV, SpVic, VPM, and S1) contain distinct topographic maps reflecting the peripheral arrangement of whiskers on the face: “barrelettes” in the brainstem, “barreloids” in thalamus, and “barrels” in cortex. However, the functional role of this somatotopic patterning, if any, remains unclear (4). Moreover, whatever the contribution of the whiskers to the appetitive component of ingestive behavior, they appear to make little or no contribution to intake and body weight regulation. In rats maintained under normal lab conditions, section of the infraorbital branch of the trigeminal nerve – which innervates the whiskers – has negligible effects upon these variables (5).

In contrast, trigeminal oral, perioral, and intraoral inputs are critical for the sensory control of eating and drinking in rodents. Deafferentation of these regions of the face is followed by a syndrome of ingestive behavior deficits including aphagia, adipsia, incisor overgrowth, impairments in the sensorimotor control of eating and drinking, and a reduction of food- or water-reinforced operant behavior. Recovered animals show a prolonged and significantly reduced responsiveness to food and water, with recovery clearly modulated by the tactile properties of the food. The reduced responsiveness is accompanied by a reduction in the level of body weight regulation to about 80% of ad lib intake (5).

However, because peripheral deafferentation abolishes sensory input equally to all three trigeminal central pathways (lemniscal, paralemniscal, and extralemniscal), it is of limited utility in identifying the specific trigeminal central pathway(s) associated with the ingestive impairments. One possible approach to more selectively dissociate those pathways is to use the *Prrxl1*^*-/-*^ “knockout” (KO). In this mutant somatotopic patterning is normal in SpVic, the start of the extralemniscal pathway, as well as the spinal caudalis nucleus (SpVc). It is selectively absent along the entire trigeminal lemniscal pathway from PrV to cortex. However, the *Prrxl1*^*-/-*^ smutation also reduces the number of primary sensory neurons in the Vg by 40-50%, a loss that affects the entire trigeminal brainstem complex. The effect is most pronounced in the PrV lemniscal nucleus, with ∼50% neural loss, compared to only ∼25% loss in the para- and extra-lemniscal nuclei of SpVi (6, 7). In this respect, the *Prrxl1*^*-/-*^mutation resembles a lemniscally-biased orosensory deafferentation.

The *Prrxl1*^*-/-*^ animal exhibits many of the deficits seen in the (recovered) peripherally deafferented rat. These include reduced eating efficiency, a reduced body weight, difficulty consuming hard food, and marked incisor overgrowth (8-10), reflecting the absence of the normal pattern of bruxism seen in rats with intact orosensory input from the incisors (11). The present study was designed to examine the ingestive behavior of this mutant at both a high temporal resolution and over extended timespans, so as to obtain baseline behavioral measures for future studies. We discuss the likelihood that the observed deficits are associated with a disruption in afference specifically along the lemniscal pathway.

## 2. Methods

All methods were approved in advance by the institutional Animal Care and Use Committee (ACUC) of Northwestern University

### 2.1. Animals

A *Prrxl*^*+/-*^ (129/B6 background) mouse was backcrossed three generations to CD1 strain. Subjects were adult wild-type (WT, *Prrxl1*^*+ / +*^) and mutant *Prrxl1*^*- / -*^ (KO) Both female and male mice littermates were used (WT 3 female, 5 male; KO 5 female, 3 male). Range of ages, in months, for both groups was similar (WT 2.3-7.6 starting, 13.6-22.6 ending; KO 4.6-7.6 starting, 12.9-21.1 ending). The *Prrxl1*^*- / -*^ mice, also known as the DRG11 line in the literature (6, 7), were generated in the Feinstein lab at Hunter College (8).

Eight WT and eight KO mice were transferred to Northwestern University from Hunter College to participate in experiments on both drinking and feeding behaviors. Mice were at least nine weeks old at the time of transfer to Northwestern and 3-20 months old during data collection. Transferred animals were housed in a reverse light cycle room, 10hr dark:14hr light, with their dark cycle starting after 8am CST. Mice from same litter were co-housed (paired) in a cage for social enrichment.

*Prrxl1*^*- / -*^ animals are fragile and require special care to survive to adulthood (10). Animals were maintained on a standard lab chow diet, supplemented as needed with Bioserv™ Nutra-Gel Diet™, a special soft gel formula that provides supplements of both food and water. Because the *Prrxl1*^*- / -*^ phenotype includes malocclusions, mice were assessed carefully at least three times weekly for signs of incisor overgrowth and the teeth were clipped when necessary. If mice displayed significant signs of distress (hunched posture, ruffled fur, low mobility, significant weight drop) they were temporarily removed from the experiment and given Nutra-Gel until their health was restored. Throughout the period of data gathering, body weights, animal appearances, animal locomotion, and total food eaten was recorded daily.

### 2.2 Experiments on drinking behavior

#### 2.2.1 Measurement of licking behavior

Mice were water deprived to a regime of 1 mL per diem, and water ingested during testing was supplemented to provide that daily amount. Testing did not start until at least seven consecutive days of water deprivation and at least 16hrs separated each daily session. All testing was done under IR illumination, with experimenters outside the room. At the start of a session the mouse was placed in a rectangular transparent acrylic elevated enclosure, 8”L×2”W×4”H, with entry/exit access only at one end.

Trials were self-initiated by the mouse leaving the enclosure and seeking the water reward. The session was terminated if the mouse either failed to leave the enclosure or did not begin licking within 15 consecutive minutes. At the start of a session a miniature (hypodermic) stainless steel water tube, coupled to a touch sensor, was positioned 1-4 inches from the entrance of the enclosure using a remotely controlled robotic arm. It was placed such that a mouse could reach the tube with their mouth but not jump on the apparatus. In order to receive a reward, the mouse had to position its head in front of the lick tube, and lick the tube. Immediately after the first lick was detected, a single water droplet (calibrated to 3 – 5 μL) was delivered. Only one droplet was delivered per trial. All mice continued to lick for some duration after droplet delivery. Between each reward, an inter-reward interval of at least 5 seconds was imposed.

The mice were free either to lick continuously during a trial and wait until the next trial or go back to the enclosure and, after some time, initiate another trial. Mice licked approximately 10 - 30 times for each reward dispensed. The robot arm holding the water dispensing system was not moved during a trial. To ensure that the mice explored the space and to avoid simply training mice on a repetitive set of movements, the tube’s position was changed every 5-10 minutes.

Water delivery was controlled by a solenoid valve and custom Arduino program and triggered on a hardware interrupt with millisecond precision. Contact data were collected with an Arduino, and timestamps of event changes recorded. Experimenters monitored this behavior in real-time through a video feed to verify drinking behavior.

Licking behavior was measured using a capacitive touch sensor (Sparkfun AT42QT1010). The sensor reported state changes and each interaction with the sensor is termed a “Contact-Detach switch” (CD), reflecting either onset or offset of touching the reward tube. Therefore, the present study does not specify lick onsets or offsets, but simply the occurrence of an interaction with the sensor, thus capturing the timepoints of the behavioral transitions. Fig. 1A is a raster plot showing all such interactions with the sensor for one mouse on one day.

**Fig 1.**
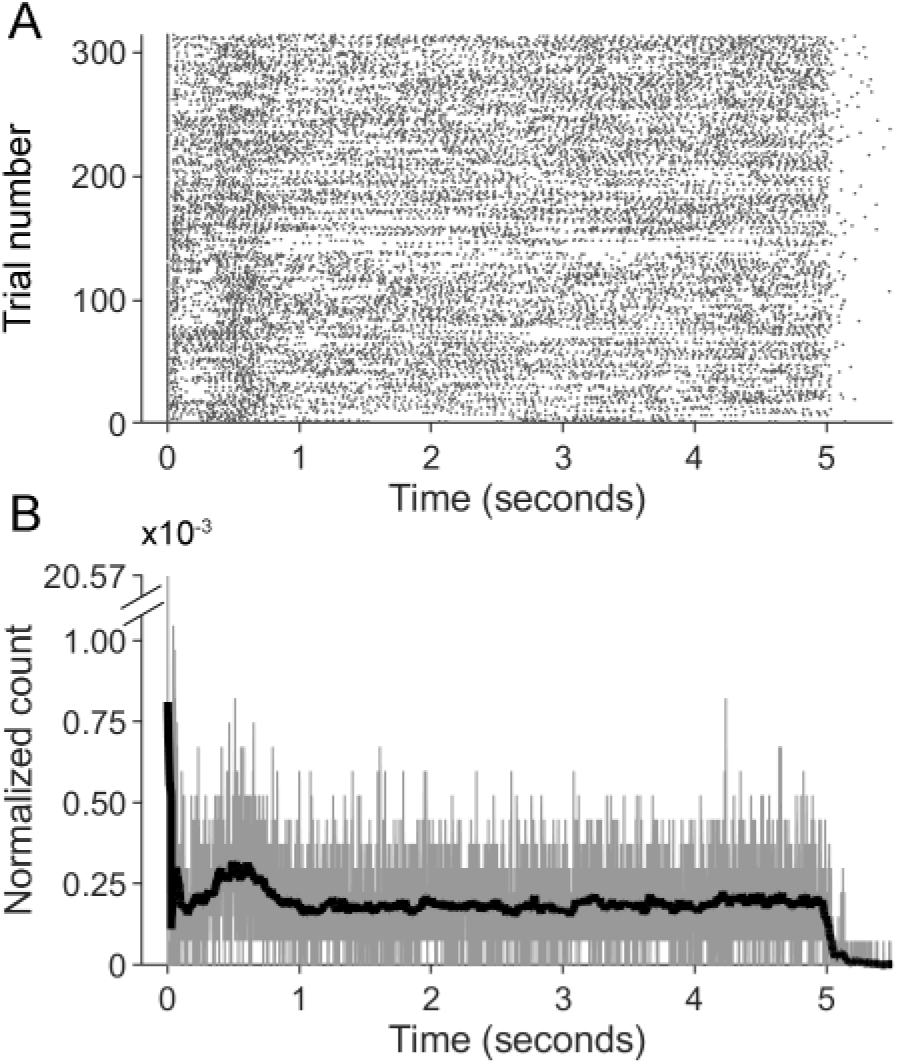
An example of measurements of sensor contact/detaches and procedure for computing histograms. (A) An example of a raster plot of contacts and detaches (CDs) from one mouse during one session. (B) To create histograms quantifying licking behavior, rasters such as that shown in (a) were summed (black trace) and then smoothed with a window of 51 msec (light gray trace, overlaid). The trials are aligned on first lick, so the large value at zero is related to the number of trials.

#### 2.2.2. Analysis of licking

The procedure for creating histograms from rasters of licking behavior is depicted in Fig. 1B. For each mouse on each day, CD (contact/detach) rasters were summed and then smoothed with a moving window of 51 msec (25 msec on either side of a central value). The value of 51 msec was chosen as a compromise: we aimed to avoid magnifying the variability between trials and between animals, while also not time-averaging over so long a duration that all temporal structure vanished. All histograms were normalized so the area under the curve was 1. Results for values of 11, 51, 81, 101, 201, and 1001 msec are shown in Supplementary Fig. S1.

### 2.3. Experiments on feeding behavior

#### 2.3.1. Monitoring food intake

Body weight and total food consumed each day (Chow pellets, Nutra-Gel) were recorded for each animal by subtracting the amount remaining from the amount provided the previous day. To ensure that mice with malocclusions had equal access to hard food, 2-6 pellets of different sizes were dispensed each day both on the cage floor and on the wire lid top.

As described in earlier studies (9, 10), *Prrxl1*^*- / -*^ animals occasionally displayed health complications, such as severe malocclusions, transient *alopecia* (particularly in overgroomed caudodorsal areas), poor locomotion, hunched posture and ruffled fur, and transient weight drops (presumably associated with insufficient consumption of hard food). During these periods, we supplemented the animals with 2-8g of soft food and/or removed them from water restriction and/or provided higher *per diem* water amounts until the symptoms resolved. All the data used in our analyses comes from animals assessed as healthy on the day of data collection.

To control for these variations in diet, the results presented in Fig. 3 include only days when animals were not receiving soft food for at least two days and were not water deprived for at least five consecutive days. Two pairs of WT animals and two pairs of KO animals were from the same litter and co-housed throughout for social enrichment. For these pair-housed individuals, we report the mean consumption per animal, that is, total amount consumed per cage divided by two. Mice were between 2 and 23 months old during the 17 months of food consumption behavior recordings.

#### 2.3.2 Assessing feeding behavior

To assess feeding behavior in frame-by-frame (30 fps) video analysis (Table 1 and Fig. 4), a mouse was placed in a 8”L×2”W×4”H acrylic tunnel-shaped enclosure with one open side. The enclosure was elevated one foot above the tabletop so that as the mouse perched on the edge of the enclosure it whisked into empty space, unless an object or reward was deliberately placed within the search space. During periods of exploration the trainer placed a “treat” (a single piece of flavored, sugared cereal (Froot Loop™) on a platform or on a tube connected to a robotic arm and then left the room. The mouse had to notice the cereal piece and reach from the tunnel to obtain the cereal. Sometimes the mouse jumped over to the platform to obtain the cereal piece, and then immediately returned to the enclosure with it. Other times the mouse stretched to the platform and grabbed the cereal without leaving the enclosure. Mice always retreated well back into the enclosure to manipulate and consume the cereal piece. A detailed analysis of these “treat trials” was carried out.

**Table 1.** Acquisition and consumption of a piece of sugared cereal quantified for WT and Prrxl1^- / -^ animals.

| Behavior | Wildtype animals | *Prrxl1*<sup>-/-</sup> animals |
| --- | --- | --- |
| No interaction with cereal piece trials | 7/27 trials | 15/30 trials |
| Interaction with cereal piece trials | 20/27 trials | 15/30 trials |
| • Knocked down cereal piece | 1/20 trials | 5/15 trials |
| • Abandoned cereal after attempting | 1/20 trials | 2/15 trials |
| • Fell while trying to obtain cereal | 1/20 trials | 1/15 trials |
| • Successfully obtained cereal piece | 17/20 trials | 7/15 trials |
| Average eating rate (g/minute) | Median : 0.063<br>Range : 0.020- 0.132 | Median : 0.012<br>Range : 0.005-0.069 |

Trials in which the mouse successfully obtained the cereal piece and began eating were deemed successful. In some trials, the mouse did not interact with the cereal at all, and on other trials, the mouse’s only interaction was to knock the cereal piece off the platform. There were also trials in which the mouse fell out of the enclosure while reaching for the cereal and in which the mouse abandoned its attempt to obtain the cereal. The numbers of all trial result types were annotated and presented separately.

For each successful trial, we manually scored the times when: 1) the mouse first interacted with the cereal piece; 2) the mouse had the cereal piece in its paws and began eating; 3) the mouse took breaks from eating; and 4) the mouse stopped eating. In the duration between timepoint (2) and time point (4), we recorded the fraction of the cereal piece, rounded to the nearest 25%, every 20 seconds.

## 3. Results

### 3.1. The Prrxl1 mutation: General description

Fig. 2 outlines the nature of the *Prrxl1*^*- / -*^ mutation as it affects somatotopy in the trigeminal system. In the WT mouse, distinct somatotopy is observed in the lemniscal pathway (PrV, VPM, and S1), as well as in SpVic and SpVc. For the *Prrxl1*^*- / -*^ deletion, somatotopy is eliminated in PrV, VPM, and SI cortex but remains intact in SpVic and SpVc. Somatotopy in the dorsal column nucleus-based lemniscal and cortical pathway were also found to be unaffected, thus the deficits in the trigeminal system associated with *Prrxl1*^*- / -*^ deletion are PrV-specific (6).

**Fig 2.**
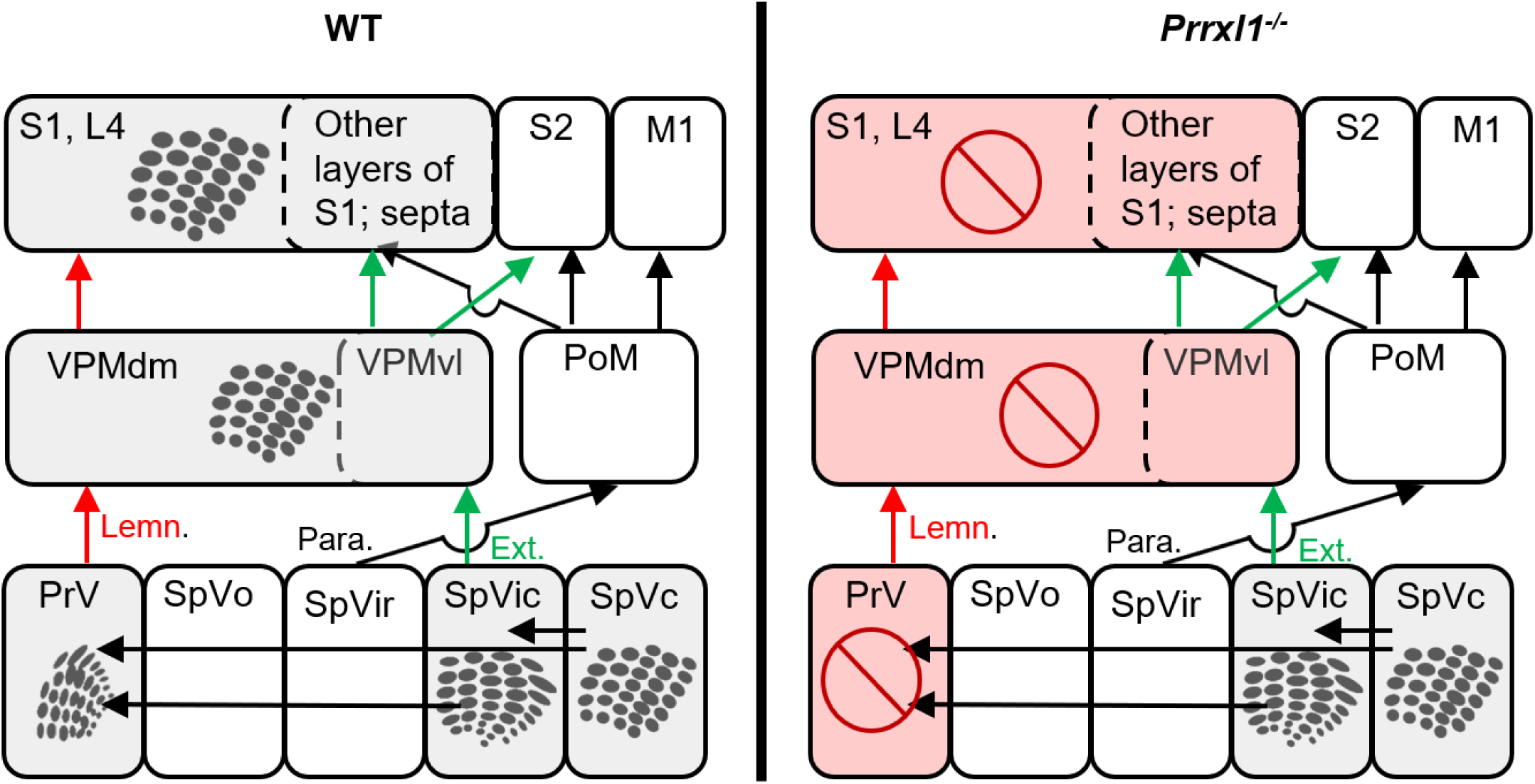
Prrxl1 deletion results in genetic ablation of somatotopy selectively along the lemniscal pathway. Somatotopic patterning in lemniscal (Lemn., red) paralemniscal (Para., black), and extralemniscal (Ext., green) pathways in WT and *Prrxl1*^*- / -*^ mice. In KO mice, somatotopy is spared in SpVic and SpVc but eliminated in PrV, VPM, and SI cortex.

*Prrxl1*^*- / -*^ animals were distinguished by a hunched posture and ungroomed fur, which made them recognizable even to untrained observers. These traits were not present on all mutant animals, and only intermittently present even for those mutants that did display them. Consistent with previous reports, we also observed that the KO animals sometimes had *alopecia* or skin lesions, which appeared intermittently and resolved over time. A smaller subset of KO animals tended to vocalize frequently (within the range audible by humans) during handling.

### 3.2. KO mice consume less hard food and maintain lower body weights than WT mice

It was sometimes necessary to provide soft food to the KO animals to maintain their body weight and resolve transient health complications (*Methods)*. To assess the animals’ ability to consume hard food pellets we selected periods when they had had *ad lib* access to water for at least 5 days, and had received no soft food for the previous 2 days.

Fig. 3 summarizes body weight and hard food consumption for WT and KO mice for these periods (sexes and ages in *Methods*). KO mice weighed significantly less than WT mice throughout the time course of 16 months that animals were housed in our facility (3a WT 42.52±7.07, Prrxl1KO 28.41±3.95; mean ± std). All but one WT mouse maintained consistently higher weight than all the KO mice (individual distributions plotted on right inset of 5a).

**Fig 3.**
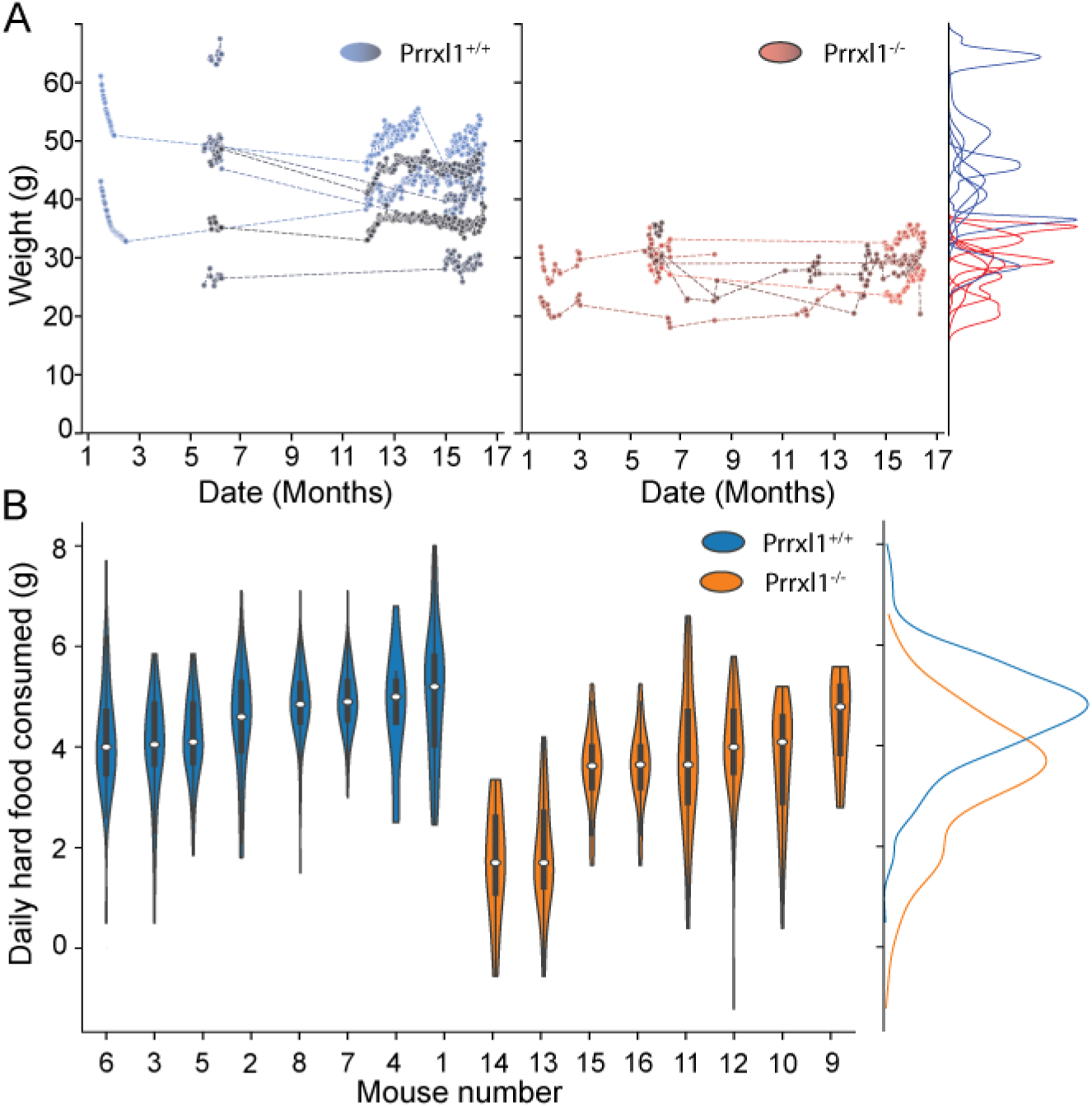
Body weights and daily food intake of WT and KO mice. Data shown for days of unrestricted access to hard food and water with at least 5 preceding consecutive ad lib water days. (A) Comparison of weights for WT and KO mice over a 17 month period. Wildtype mice (left panel, dark blue to black) consistently maintain higher body weights than KO mice (right panel, light red to brown). Only one WT animal overlaps with the KO population (individual distributions as inset on far right. WT=blue, KO=red) (B) KO mice (orange distributions, right side) consumed on average less hard food than wildtype mice (blue distributions, left side): WT 4.66±1.03 g, Prrxl1KO 3.41± 0.32 g; mean ± std, p=6.83e-38 Welch’s test. 6 out of 8 KO animals have means equal to or lower than the lowest WT. Interquartile ranges plotted as dark bars. Population distributions plotted as inset on the far right using the same color scheme.

Because the weight data suggested that KO animals were eating less food we next quantified the amount of hard food consumed on each day (3b) The weight of the hard pellets present in the cage on a given day was subtracted from the weight of the food placed in the cage on the previous day. On average, WT mice consumed slightly more hard food (WT 4.66±1.03 grams, KO 3.41± 0.32 grams; mean ± std, p=6.83e-38 Welch’s test). 6 out of 8 KO mice ate less, on average, than the WT mouse with the lowest average consumption. However, the distributions for KO and WT animals overlap substantially, a puzzling result given the large discrepancy in body weights. One possible explanation is that KO mice may exhibit inefficient eating behavior, such that when they bite the hard food pellets, a portion is lost on the cage floor as small fragments that could not be measured in the present experiment. An inefficient “sloppy eater” phenotype was previously observed in *Prrxl1*^*-/-*^animals that were reared and maintained on a liquid diet (8). We next sought to quantify their eating efficiency for sweet hard food (sugared cereal), which is strongly preferred by the mice.

### 3.3. Prrxl1^-/-^mice are less efficient and less precise in their eating behavior

To measure eating efficiency, mice were presented with a single piece of sugared cereal, in a dark room, with the experimenter absent and their localization and consumption behaviors quantified. Mice had to perch from a housing enclosure, localize the cereal placed on an elevated platform with a gap between the enclosure and the platform, and grasp the cereal with their mouth (see *Methods*). Table 1 summarizes our findings.

The oral grasping behavior of KO mice was infrequent and inefficient compared to wildtype animals. KO animals interacted with the cereal on only 50% of the trials, compared to 75% for WT animals. On those trials in which the animals did interact with the food, all but one of WT animals (7/8; 88%) successfully grasped the cereal on nearly all (17/20; 85%) trials. In contrast, less than half of the animals (3/8; 38%) were successful in grasping. In addition, those KO animals that were successful in grasping, were successful on less than half of the trials (7/15; 47%). The remaining 5/8 KO animals either did not interact with the food (2/5 animals) or did not successfully grasp it (3/5) to start an eating session. KO mice often bumped into the cereal and knocked it off the stage or abandoned attempts to eat.

No difference between the genotypes was found in the time it took the animal to find the food, because this “discovery duration” depended on what the mouse happened to be doing at the time of its presentation. If the mouse was already at the edge of the tunnel, then it found the cereal rapidly (within 1 – 2 seconds), whereas if it was turned away from the tunnel entrance then it took much longer to find the cereal (several minutes). Similarly, no differences were observed in the number of approaches that WT and KO mice made towards the food (usually 1-3 approaches for both genotypes). Approaches were scored as times when the mice moved with their head or body oriented towards the food, whether or not they continued to interact with the food.

We next analyzed the minute-by-minute ingestion behavior for all 7 KO trials and 15/17 WT trials with successful cereal grasping (Fig. 4). Note, this analysis necessarily involves a small number of trials, since the mutants were so unsuccessful in obtaining the “treat”. Two of the WT trials (mouse 7) were not analyzed because the animal knocked the cereal piece off the platform after it had started eating. Two of the three KO mice that obtained and consumed the cereal displayed eating durations more than twice as long as any of the WT mice. One of the KO mice was unable to finish more than half of the cereal, in all four trials. Only one of the three KO mouse was successful in rapidly eating the entire cereal in the two trials where they tried. In contrast, all WT mice showed fast and efficient eating behavior: on only 2/17 trials did a WT mouse stop eating the cereal part ways. KO mice eating rate in all trials for which food weight data was available (6/7 KO trials, including the two fast eating trials, 13/15 WT trials) was substantially lower than that of WT animals (Table 1, last row).

**Fig 4.**
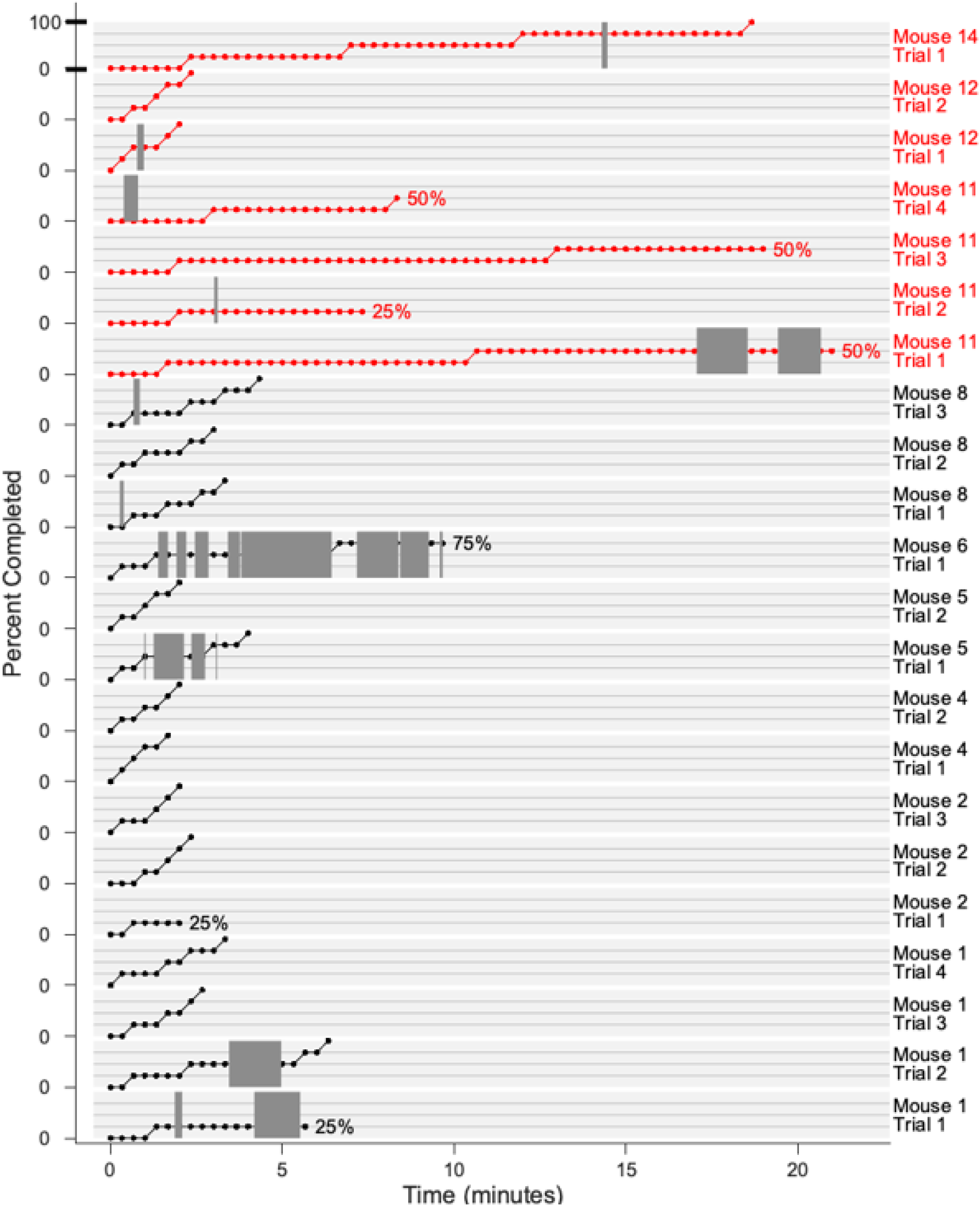
Consumption of a piece of sugared cereal as a function of time. Each light gray horizontal rectangle indicates a separate trial, with mouse number and trial number indicated on the far right. The vertical axis for each rectangle ranges between 0% and 100%, as indicated for the top plot. The 0% location is marked on the vertical axis for each trial separately. Horizontal grid lines across each rectangle indicate the 25%, 50%, and 75% levels. Data points are located every 20 seconds (dots). Within each trial, dark gray regions indicate times during which mice took breaks from eating.

In summary, a detailed comparison of the ingestive behavior of WT and KO mice, at sub-second temporal resolution, indicates that *Prrxl1*^*- / -*^ mice tend to be less efficient in oral grasping of food and modulating oromotor sequences for eating.

### 3.4. KO animals exhibit less consistent and less persistent licking behavior than WT animals

When water was delivered through a reward spout, both WT and KO animals licked approximately 10-30 times for each reward dispensed. Examples of typical contact-detach (CD) rasters for a WT and KO mouse are shown in Fig. 5A. In this example, the WT mouse (mouse 4) generated 69, 100, and 122 drinking trials during its first three sessions, respectively. For three equivalent sessions, a KO mouse (mouse 11, red right panel) initiated 69, 67, and 89 trials. Given that the mice had equal opportunities for licking, these data suggest a reduced responsiveness to water in the KO mouse. Inspection of Fig. 5A also suggests a reduction in the persistence of such interactions in the KO mouse, as reflected in the reduced density of the tick marks in the raster. Finally, the raster suggests higher variability in the KO behavior and less trial-to-trial consistency: later trials in each session have a lower density of events than trials earlier in a session.

**Fig 5.**
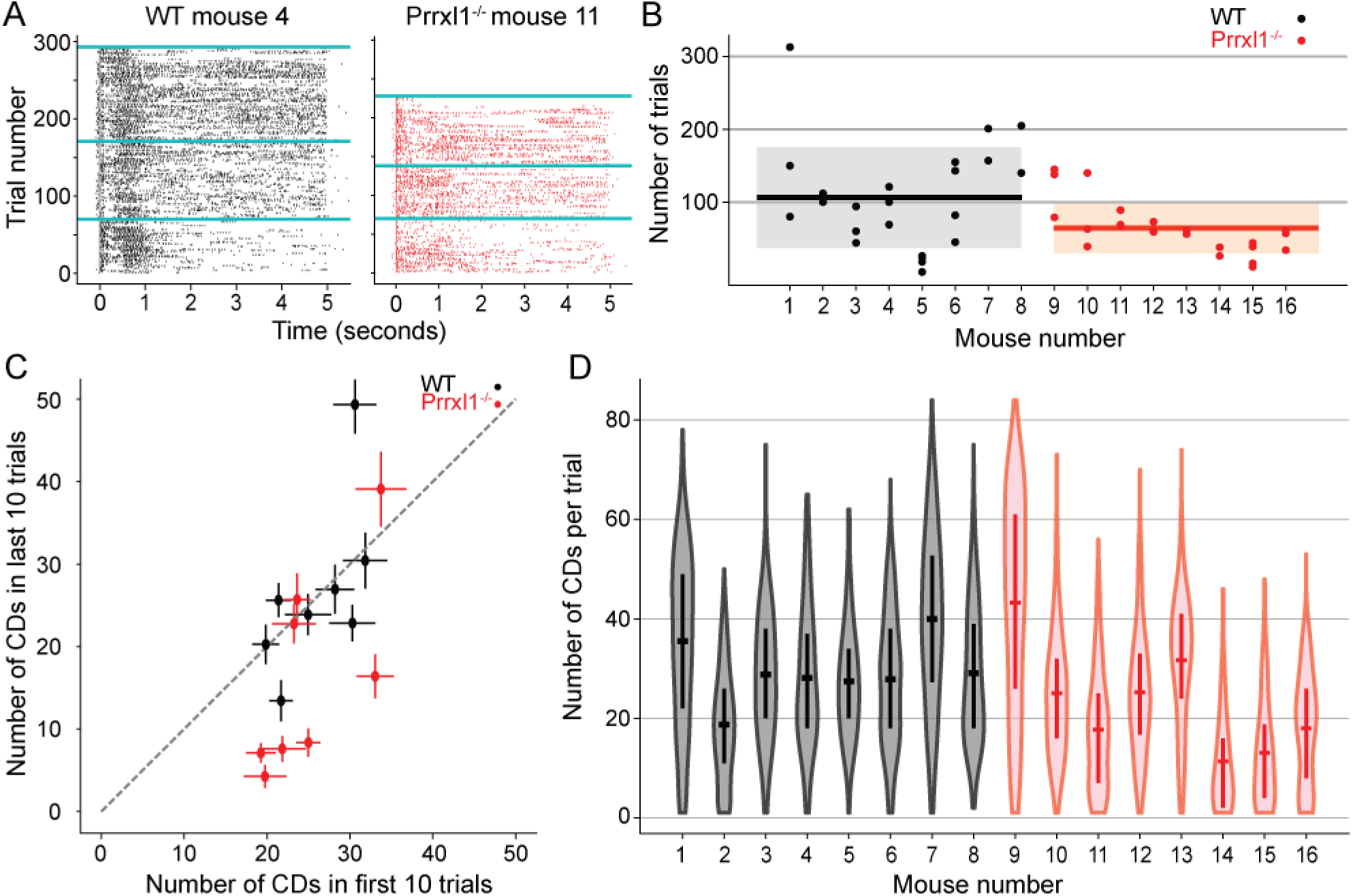
Knockout animals are less persistent and less consistent in their drinking behavior. Comparison of number of trials performed per experimental session, and number of CDs per trial for WT and KO mutant mice. (A) Typical CD raster plots WT (mouse 4, left) and KO (mouse 11, right). Each tick represents either a contact or a detach from the lick sensor. Both mice participated in three experimental sessions; sessions are divided by cyan horizontal lines. (B) WT (mice 1-8, black) tended to perform more trials than KO (mice 9-16, red). Means for both cohorts are shown as black horizontal lines, with standard deviations indicated by semi-transparent gray and red rectangles, respectively. (C) KO mice were less consistent in licking during a trial than the WT mice. Mouse numbers and colors as in (B). Licking activity for five of the eight KO mice falls well below the diagonal, while WT mice cluster close to the equality line. (D) This plot controls for the possibility that KO animals have a higher lick rate than WT and thus fatigue more quickly. KO animals tend to perform slightly fewer CD/trial on average (mean ± ste: 27.2 ± 0.5 CDs/trial for KO versus 30.4 ± 0.3 CDs/trial for WT; p < 1e-8, two-sided t-test), thus the difference in the number of trials per session (B) and the decreased lick rate towards the end of the session (C) are not explained by a difference in effort spent per trial. Mouse numbers and colors as in (B).

Fig. 5B-C generalize and quantify these results over all mice. The data plotted in Fig. 5B show that individual mice varied greatly in the number of trials generated during each session. KO mice tended to perform significantly fewer trials than WT (mean ± std: 64 ± 36 trials/session versus 107 ± 71 trials/session; p = 0.012, two-sided t-test). Notably, four of the WT mice (mice 1, 6, 7, and 8) had one or more sessions in which they performed 150 trials or more, while none of the KO mice ever performed more than 145 trials per session.

Fig. 5C indicates that KO mice were also less consistent in their licking responses. The number of CDs in the first 10 trials of each session did not differ between KO and WT groups, suggesting a similar level of initial thirst. However, 5/8 KO mice generated fewer CDs in the last 10 trials of each session compared to the first 10 trials. Four of those five animals exhibited a ∼67% reduction. In contrast, only two of the eight WT animals showed a reduction in the number of CDs in the last 10 trials compared to the first 10 trials. Moreover, that reduction was less severe (only ∼50% of the starting trials). Taken together, results in 3b-c suggest a systemic reduction in responsiveness to water in KO animals, replicating the results of deafferentation studies.

An alternative explanation for the data of Fig. 5C is that KO mice could have increased their licking rate mid-session and then fatigued earlier, thus generating fewer total trials as well as fewer CDs at the end of the session. However, when we compare the entire distribution of CDs per trial for each animal in Fig. 5D, the two populations overlap. Moreover, they differ in a direction opposite to that which would be predicted from a higher mid-session licking response in KO animals (mean ± ste: 27.2 ± 0.5 CDs/trial for KO versus 30.4 ± 0.3 CDs/trial for WT; p < 1e-8, two-sided t-test), with 3/8 KO mice showing a smaller mean CD/trial from any of the WT animals.

### 3.5 Relation between eating and drinking behaviors

Because it is well-known that food and water intake are correlated (12), we next show a comparison of the daily food consumption data and the number of lick trials (Fig. 6). Within a genotype, animals that performed many licking trials also tended to consume more daily food, and conversely, animals that tended to eat less daily also performed fewer lick trials. In addition, the ingestive data space suggests a functional division between the two genotypes, with Prrxl1^-/-^occupying a distinct, but partially overlapping region with wildtype mice. With the previous findings suggesting a sensory driven ingestive deficit, we next examine the temporal profiles of the licking behavior time course.

**Fig 6.**
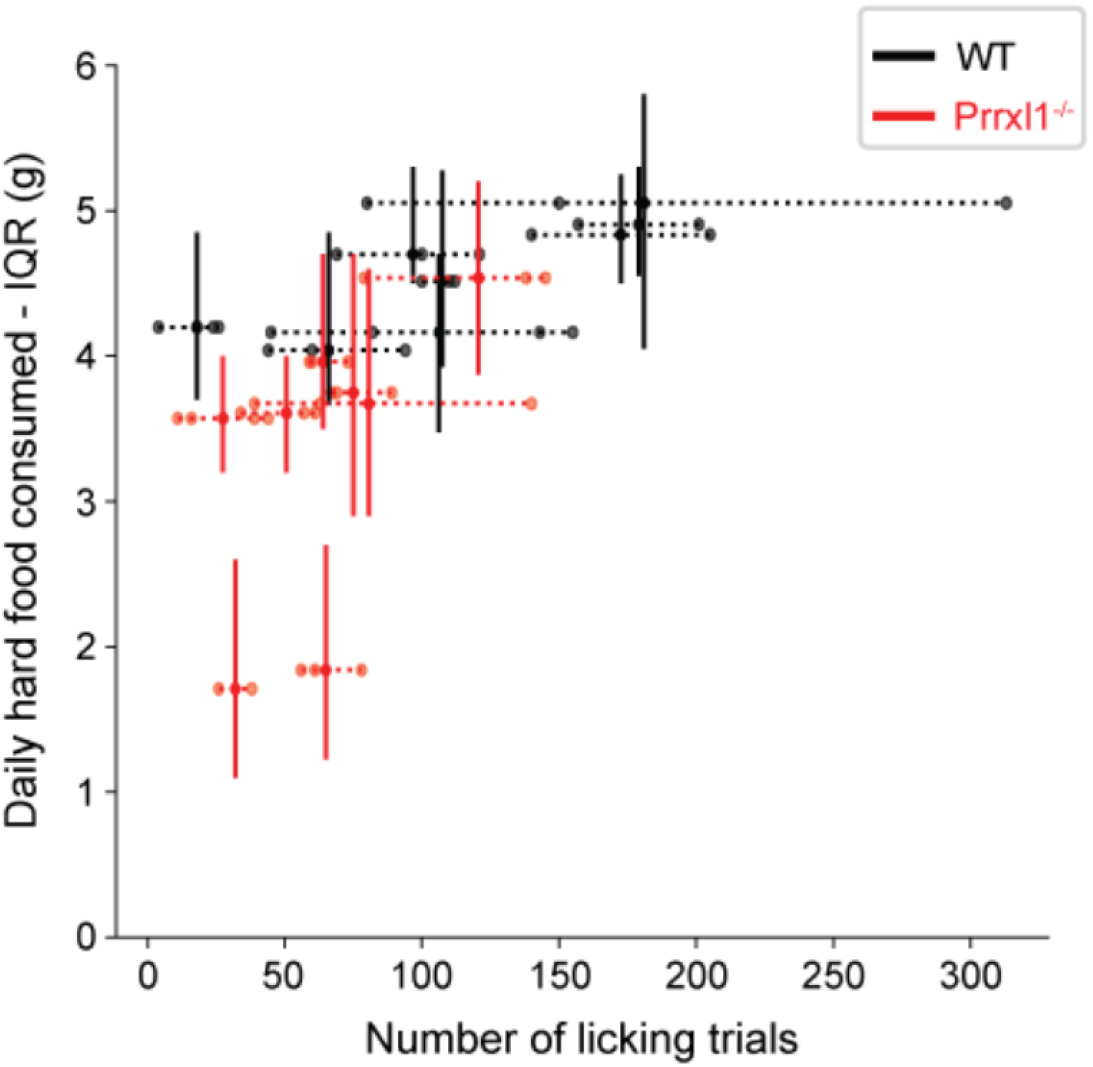
A comparison of food consumption and the number of licking trials. Within a genotype, animals that performed more licking trials also ate more and animals that ate less tended to perform fewer licking trials. Two of the eight mutant animals (#13 and #14) display more severe food intake issues. Both groups occupy separate regions that partially overlap, suggesting two functionally distinct group in this projection of the ingestive behavior space.

### 3.6. WT and KO mice differ significantly in the time course of licking and ability to modulate licking rate

Fig. 7 compares the time course of licking for WT and KO mice populations as reflected in histograms generated from the raster plots (see *Methods*). WT mice (7A, black/grey traces) exhibited a pronounced modulation of licking rate upon sensory contact. This change in the lick rate starts a few milliseconds after first contact, when water delivery occurs, and peaks 100-1000 msec after reward delivery. The population histogram for all KO mice (7B) displays considerably more variability, with some trials showing a decrease in licking between 100-1000 msec, followed by its resumption near 1000 msec. The standard deviation of the CD histogram shown in Fig. 7B is particularly broad between 100-1000 msec, suggesting that after their initial contact with the water tube, KO mice are differentially delayed at initiating repetitive licking motions, and there is larger inter-trial or inter-animal variability in this consummatory time window. In addition, the variability appears bimodal and symmetric, so that the mean of the KO CD histogram is essentially flat. The inset to panel 7B overlays the mean liking rates for WT and KO animals, revealing one of the strongest differences in orosensation observed.

To more closely examine the bimodal variability across the KO mice, Fig. 7D shows the time course of licking behavior for each mouse in order to assess the difference and consistency in the amount of lick modulation between the two groups. All WT mice modulate their licking response, precisely and rapidly upon sensory contact. In contrast, only two out of the eight KO mice (animals 9 and 13) show modest modulation of their licking behavior (8/8 WT versus 2/8 KO; p = 0.007, Fisher’s exact test). The modulation is significantly delayed, starting after 500ms and peaking around 1s after water reward onset.

**Fig 7.**
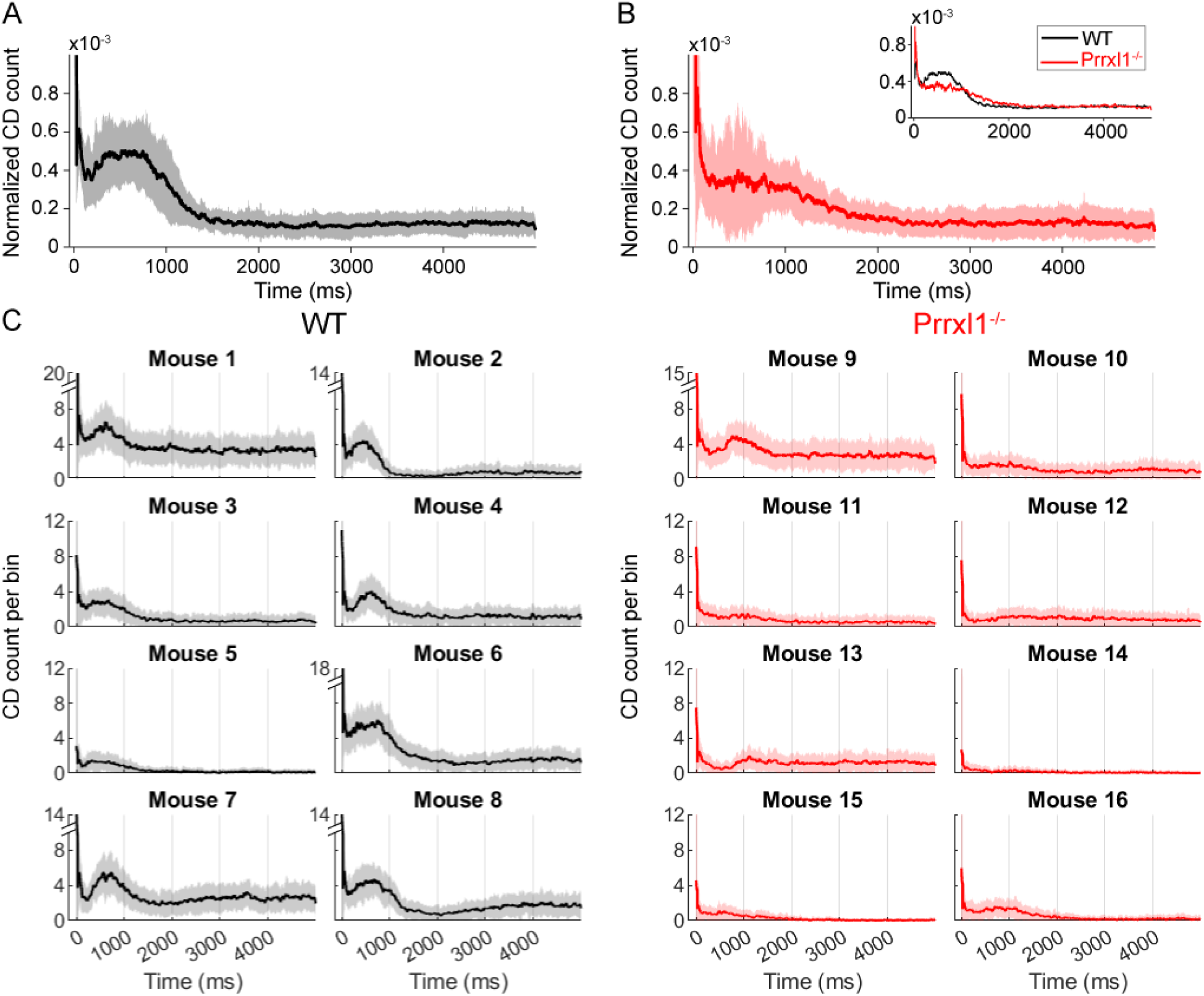
Time course of licking in WT (black and gray) and KO (red and pink) mice. (A) All wildtype mice showed a peak in licking behavior between 1-1000 msec after the water reward was dispensed. The black curve shows the mean of the CD histogram computed across all sessions and all days for all wildtype mice. Plus and minus standard deviation around the mean is shaded gray. (B) *Prrxl1*^*–/–*^ mice showed much more variable licking behavior. The red curve shows the mean of the CD histogram computed across all sessions and all days for all *Prrxl1*^*–/–*^ mice, with plus and minus standard deviation around the mean shaded light red. The *inset* overlays the mean CD counts for WT and KO animals (C) All individual mice CD histograms with a bin size of 51ms. All y-axes go to 12 CD counts/bin unless labeled with an axis break, in which case the maximum y-value is labeled.

## 4. Discussion

In the present study we examine food and water intake and the temporal organization of eating and drinking movements in the Prrxl1^- / -^ mutant mouse. With appropriate husbandry (10), this mutant can initiate drinking, eat hard food, and live an extended life. This is an important consideration in view of its unique contribution as a model system for the study of such problems as pain, ingestive behaviors, and the origin and significance of somatotopic patterning in the mammalian somatosensory system.

### 4.1. Trigeminal central pathways and the control of feeding behavior in rodents

Deafferentation studies (e.g., (5), reviewed in (13)) have clearly identified trigeminal afference as a key source of the peripheral feedback signals driving ingestive behavior in rodents. The data in the present study were obtained in a preparation quite different from deafferented rats, and also quite different from the *Prrxl1*^*-/-*^KO described in (8). In neither of these earlier studies could the animals generate the ingestive behaviors required to maintain themselves on hard food. In contrast, the present mice, through a combination of improved husbandry and outbreeding to the CD1 mouse strain, could generate well-organized ingestive behavior sequences. In this respect, they resemble recovered deafferented animals, who could sustain themselves on hard food (5). The behavior of both the deafferented rats and the *Prrxl1*^*- / -*^ mice reflects an adjustment to chronic afferent disruption, but on two different time scales: short-term, during recovery, for the deafferented rats; long-term, from birth, in the mutant. The *Prrxl1*^*-/-*^mutant thus allows the study of ingestive behavior in an animal whose ingestive motor sequences are present, but whose trigeminal central pathways are genetically perturbed throughout its life. *Prrxl1*^*-/-*^

The present work shows that *Prrxl1*^*- / -*^ animals can perch from an enclosure and use their tactile senses in complete darkness to find and subsequently consume water from a water spout whose position in space was randomly varied over time. They could also lick this spout to initiate a water reward. They did not differ from WT in the amount of licking elicited by a water reward. Importantly, they were able to ingest hard food and were viable, though at significantly reduced body weights than WT mice. Where they most resembled the deafferented rats or early mutant mice was in the reduced efficiency of their ingestive behavior This was most dramatically shown in the increase in unsuccessful grasps and in the increased time taken to consume a given unit of food.

### 4.2. Trigeminal modulation of oromotor sequences operates across several timescales

Eating and drinking are guided by a continuous stream of sensory information. While olfaction and vision provide distance information for localizing nutrient sources, successful ingestion depends upon a continuous assessment of the texture, hardness, temperature, and other mechanosensory properties of a food which guide the initial grasping and subsequent intraoral manipulation of the food source. All these behaviors are mediated by the jaw motor system (which is intact in the mutant) so that their impairment suggests a break in the flow of orosensory, primarily trigeminal, mechanosensory afference which elicits and modulates licking, grasping, chewing, and intraoral manipulation behaviors. This disruption operates over multiple timescales.

On a reflex timescale, an earlier study (14) showed (1) that the jaw opening elicited by perioral contact of the face with a food pellet or sipper tube in the normal rat was either abolished or significantly delayed in peri-orally deafferented rats; (2) that mechanical or electrical stimulation of the orofacial region elicited reflex activity in the motoneurons of the jaw-opener muscle; and, (3) that the most effective sites for eliciting activity in the jaw-openers appeared to cluster about the region of the upper lip and superior portions of the oral cavity. Indeed, for this region, mechanical displacement of less than 1 g with a Von Frey hair was often sufficient to elicit jaw motoneuron activity in these anesthetized preparations. These observations suggest that oral and perioral regions originate the afferent component of trigeminal sensorimotor circuitry that monitors the presence and location of food and water sources and that provides continuous, moment-to moment-feedback during normal licking, grasping and intraoral manipulation. Furthermore, input from the teeth also generates the bruxism which, in normal rodents, prevents the development of malocclusion--a defining phenotype of the *Prrxl1*^*-/-*^mutant.

The role of trigeminal afference in modulating either the initiation of, or ongoing, ingestive behaviors extends to longer chronic timescales in both “recovered” deafferented and *Prrxl1*^*-/-*^preparations. The reduction in weight and consumption rates reflects an adaptive adjustment to chronically reduced sensory input, and confirms the contribution of trigeminal afference in modulating responsiveness to food (5). These results link trigeminal orosensation to internal states of hunger and appetitive control (15); e.g., hedonic salience (16).

The effects of reduced trigeminal afference on the modulation of ongoing oromotor behaviors is especially striking at intermediate timescales, on the order of 100’s of ms. Indeed, the striking contrast between the precise and rapid modulation of licking rates observed in the wildtype animals and its delayed or complete absence in *Prrxl1*^*- / -*^ suggests a disrupted sensory-motor loop connecting trigeminal inputs to oromotor circuits. The timing of this behavior, which is delayed by hundreds of milliseconds in the mutant, suggests that in the mutants we are not dealing simply with the effects at a reflex level but with the involvement of higher order circuits. There is substantial top-down innervation of trigeminal sensory nuclei (17-20), and this input modulates *in vivo* neural responses (21, 22). In addition, decorticate preparations have suggested a modulatory role for higher order structures in ingestive behaviors (23). The relative contributions of cortical vs brainstem structures to the trigeminal control of ingestive behaviors are important questions for future research in this mutant.

### 4.3. An hypothesized selective role for the lemniscal pathway modulating ongoing ingestive behavior

The impairments described in this report have two possible explanations. First, they could result solely from the overall reduced cell number in Vg, SpVi, and PrV. The reduced cell number would in turn reduce signal fidelity in any or all of these structures. Alternatively, the ingestive impairments could be a result of a selective disruption in trigeminal afference along the lemniscal pathway.

Support for this hypothesis comes from the study by Ding, et al (6) of the impact of Prrxl1 deletion on the lemniscal pathway. First, *Prrxl1* is not expressed in SpVi (6, 24) or in lamina III or IV or SpVc, where barrelettes develop. It is expressed in PrV (where barrelettes are abolished with the gene deletion) and lamina I and II of SpVc (layers that don’t include barrelettes). Correspondingly, barrelette patterning is normal in SpVic (the extralemniscal brainstem nucleus) and SpVc, but absent in PrV (the lemniscal nucleus). Second, the absence of this patterning is associated with a reorganized axonal projection pattern at multiple stages along the lemniscal pathway. Not only do afferents to PrV fail to organize into clear whisker-specific clusters, but thalamic inputs to layer IV S1 cortex distribute uniformly instead of organizing into barrel-sized clusters as observed in WT animals (6). Lesion studies have shown that the origin of this sensory map disorganization throughout the lemniscal pathway must lie in PrV itself, and not in thalamus or cortex, where *Prrxl1* is not expressed (25).

Given the well-known feedback projections from S1 to PrV (18-20), disorganization of these thalamocortical inputs seems likely to contribute to the behavioral disruptions at the intermediate timescale. Disordered sensory organization may cause temporally-jittered, or noisy, flow of trigeminal information leading to the impaired modulation of ingestive behavior. A selective role for the lemniscal pathway is also suggested by the observation that the ingestive impairments of the *Prrxl1*^*- / -*^ animals included not only problems with the oral grasping manipulation of food objects, but with their initial localization (Table 1), suggesting some disruption in vibrissal sensory localization function. The extent to which altered circuit properties and temporal jitter affect the *Prrxl1*^*- / -*^ mutant will require electrophysiological studies comparing the response properties of SpVi and PrV neurons in wildtype and mutant animals.

*Prrxl1* is also expressed in the geniculate ganglion (GG) (24). This sensory ganglion receives gustatory information from the anterior two-thirds of the tongue, as well as mechanosensory input from the outer ear (pinna). *Prrxl1* expression in the GG thus raises the possibility that gustatory afferents might be affected by its deletion. However, a recent transcriptomic study showed that the neurons in GG that express Prrxl1 are those that receive mechanosensory input from the pinna (26). This result rules out disruptions in lingual afference or gustation as the main mechanisms for the observed deficits.

Recent findings on vibrissal tactile sensing have suggested that the lemniscal pathway may make a unique contribution to sensory coding in the trigeminal system. Yu Et al. (3) and Moore Et al. (27) showed that neurons in the trigeminal lemniscal, but not the paralemniscal pathway are “substantially modulated” by both touch and self-motion, a coding property which is likely to be critical for whisker-mediated discrimination. Chakrabarti and Schwarz (21) recorded from both PrV and SpVi neurons to show that sensory gating during an active whisking task affects the lemniscal, but not the extralemniscal, processing stream, and that modulatory input comes from sensorimotor cortex.

In summary, our results indicate that the ordered assembly of trigeminal sensory information is critical for the rapid and precise modulation of motor circuits driving ingestive action sequences. This trigeminal modulation is observed at multiple timescales, from milliseconds, to minutes, to months, tightly linking somatosensation and ingestion, from moment-to-moment consummatory to long term appetitive control. Our data also suggest that the lemniscal component of the ascending trigeminal pathway makes a selective contribution to that process.

## Supporting information

**Fig S1.**
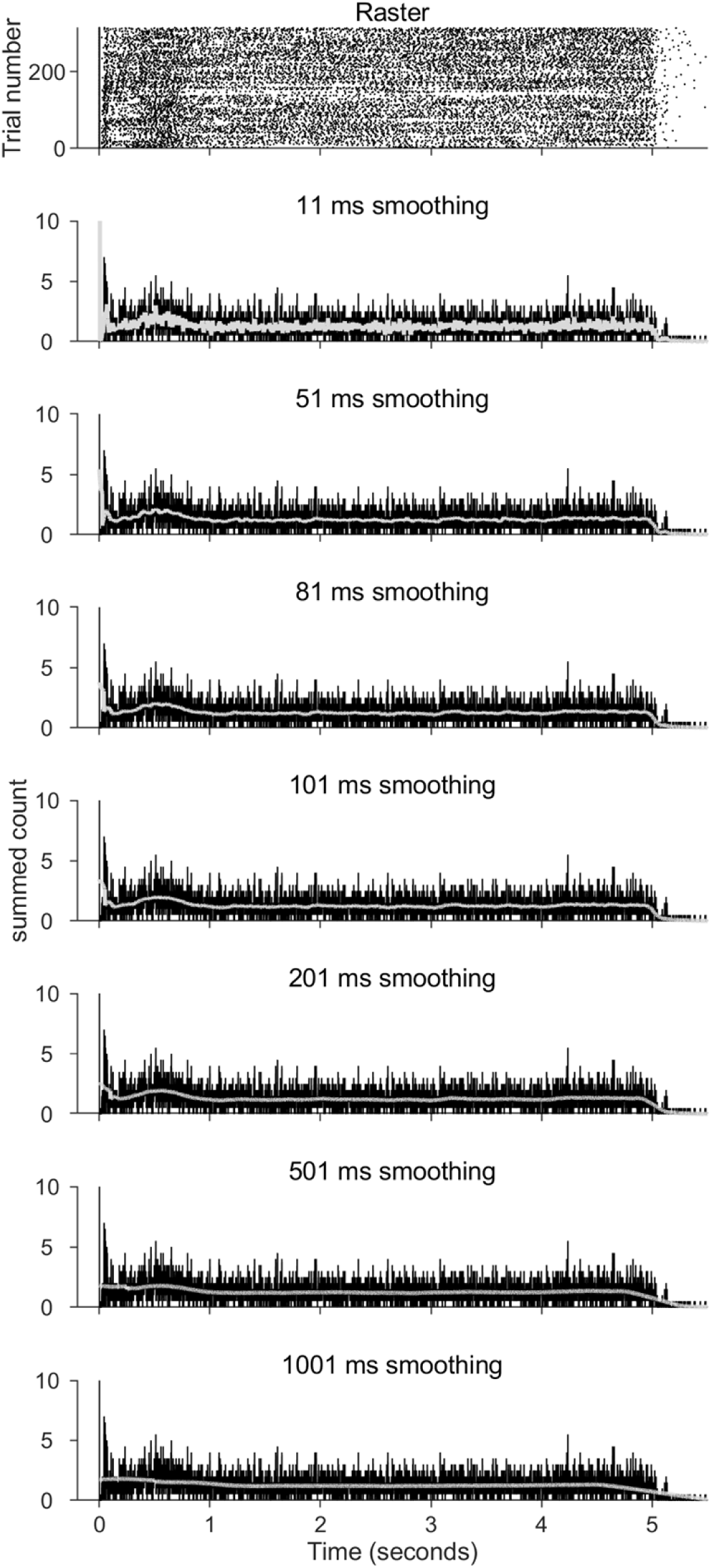
Licking contacts and detaches (CDs) shown for different smoothing windows. An example of a raster plot of contacts and detaches (CDs) from one mouse during one session shown in top panel. Effect of smoothing the summed histogram from the raster data on the top panel shown for window sizes of 11, 51, 81, 101, 201, and 1001 ms in respectively labeled panels. Black traces: summed raster data in 1ms bins, same for all smoothed histograms and overlaid for comparison. Light gray traces: the temporally smoothed average. Based on these temporal averaged data, 51ms time window chosen as the window size that preserves the temporal profile, and averages out the intertrial noise. The trials are aligned on first contact, so the large value at zero is related to the number of trials.

## Acknowledgments

The authors gratefully acknowledge the assistance of undergraduates Nadia Khan and Allison O’Donnell for help with the behavioral licking data collection. Undergraduates Olivia Asmar, Olivia Cong, Nadia Khan and Allison O’Donnell are acknowledged for help with collecting eating video data. In addition, we thankfully acknowledge all the following undergraduates for their help maintaining the animal colony and recording daily weight and food data: Olivia Asmar, Olivia Cong, Nadia Khan, Allison O’Donnell, Sanjana Sharma, William Zeng and Kevin Zhang. We thank Research Engineer Kevin Kleczka and undergraduate Gretchen Vogt for help in design and construction of components of the experimental setup. We also thank and acknowledge Sergio Bernal-Garcia for many useful discussions and for helping to transfer the *Prrxl1* animals to our laboratory. Lastly, we are very grateful to Clinical Veterinarian Rebecca Erickson at Northwestern University for frequent visits and clinical care for our *Prrxl1* animal colony.

## Author Contributor Roles

Admir Resulaj: Conceptualization, Data Curation, Formal Analysis, Investigation, Methodology, Software, Supervision, Visualization, Writing – Original Draft Preparation, Writing – Review & Editing

Jeannette Wu: Data Curation, Formal Analysis, Investigation, Methodology, Validation, Visualization, Writing – Original Draft Preparation; Writing – Review & Editing

Mitra J. Z. Hartmann: Conceptualization, Data Curation, Formal Analysis, Funding Acquisition, Methodology, Project Administration, Software, Supervision, Validation, Visualization, Writing – Original Draft Preparation; Writing – Review & Editing

Paul Feinstein: Conceptualization, Methodology, Visualization, Writing – Review & Editing

H. Phillip Zeigler: Conceptualization, Funding Acquisition, Supervision, Writing – Original Draft Preparation; Writing – Review & Editing

## References

1. El-Boustani S, Sermet BS, Foustoukos G, Oram TB, Yizhar O, Petersen CCH. Anatomically and functionally distinct thalamocortical inputs to primary and secondary mouse whisker somatosensory cortices. Nature Communications. 2020;11(1).

2. Pierret T, Lavallee P, Deschenes M. Parallel streams for the relay of vibrissal information through thalamic barreloids. Journal of Neuroscience. 2000;20(19):7455–62.

3. Yu CX, Derdikman D, Haidarliu S, Ahissar E. Parallel thalamic pathways for whisking and touch signals in the rat. Plos Biology. 2006;4(5):819–25.

4. Kaas JH. Topographic maps are fundamental to sensory processing. Brain Research Bulletin. 1997;44(2):107–12.

5. Jacquin MF, Zeigler HP. Trigeminal orosensation and ingestive behavior in the rat. Behav Neurosci. 1983;97(1):62–97.

6. Ding YQ, Yin J, Xu HM, Jacquin MF, Chen ZF. Formation of whisker-related principal sensory nucleus-based lemniscal pathway requires a paired homeodomain transcription factor, Drg11. Journal of Neuroscience. 2003;23(19):7246–54.

7. Jacquin MF, Arends JJA, Xiang CX, Shapiro LA, Ribak CE, Chen ZF. In DRG11 knock-out mice, trigeminal cell death is extensive and does not account for failed brainstem patterning. Journal of Neuroscience. 2008;28(14):3577–85.

8. Bakalar D, Tamaiev J, Zeigler HP, Feinstein P. Abolition of lemniscal barrellette patterning in Prrxl1 knockout mice: Effects upon ingestive behavior. Somatosensory and Motor Research. 2015;32(4):236–48.

9. Monteiro C, Cardoso-Cruz H, Matos M, Dourado M, Lima D, Galhardo V. Increased fronto-hippocampal connectivity in the Prrxl1 knockout mouse model of congenital hypoalgesia. Pain. 2016;157(9):2045–56.

10. Monteiro C, Dourado M, Matos M, Duarte I, Lamas S, Galhardo V, et al. Critical care and survival of fragile animals: The case of Prrxl1 knockout mice. Applied Animal Behaviour Science. 2014;158:86–94.

11. Wang CZ, Shi M, Yang LL, Yang RQ, Luo ZG, Jacquin MF, et al. Development of the mesencephalic trigeminal nucleus requires a paired homeodomain transcription factor, Drg11. Molecular and Cellular Neuroscience. 2007;35(2):368–76.

12. Possidente B, Birnbaum S. Circadian rhythms for food and water consumption in the mouse, Mus musculus. Physiology & Behavior. 1979;22(4):657–60.

13. Zeigler HP, Jacquin MF, Miller MG. Trigeminal orosensation and ingestive behavior in the rat. Prog Psychobiol Physiol Psychol. 1985;11:63–196.

14. Zeigler HP, Semba K, Jacquin MF. Trigeminal reflexes and ingestive behavior in the rat. Behavioral Neuroscience. 1984;98(6):1023–38.

15. Zeigler HP. Brainstem orosensorimotor mechanisms and the neural control of ingestive behaviour. Appetite: neural and behavioural bases New York: Oxford University Press p. 1994:54-85.

16. Berridge KC, Kringelbach ML. Affective neuroscience of pleasure: reward in humans and animals. Psychopharmacology. 2008;199(3):457–80.

17. Jacquin MF, Wiegand MR, Renehan WE. Structure-function relationships in rat brain stem subnucleus interpolaris. VIII. Cortical inputs. Journal of Neurophysiology. 1990;64(1):3–27.

18. Killackey HP, Koralek K-A, Chiaia NL, Rhoades RW. Laminar and areal differences in the origin of the subcortical projection neurons of the rat somatosensory cortex. Journal of Comparative Neurology. 1989;282(3):428–45.

19. Smith JB, Watson GDR, Alloway KD, Schwarz C, Chakrabarti S. Corticofugal projection patterns of whisker sensorimotor cortex to the sensory trigeminal nuclei. Frontiers in Neural Circuits. 2015;9.

20. Wise SP, Jones EG. Cells of origin and terminal distribution of descending projections of the rat somatic sensory cortex. Journal of Comparative Neurology. 1977;175(2):129–57.

21. Chakrabarti S, Schwarz C. Cortical modulation of sensory flow during active touch in the rat whisker system. Nature Communications. 2018;9.

22. Furuta T, Urbain N, Kaneko T, Deschenes M. Corticofugal Control of Vibrissa-Sensitive Neurons in the Interpolaris Nucleus of the Trigeminal Complex. Journal of Neuroscience. 2010;30(5):1832–8.

23. Whishaw IQ, Schallert T, Kolb B. An analysis of feeding and sensorimotor abilities of rats after decortication. Journal of Comparative and Physiological Psychology. 1981;95(1):85–103.

24. Rebelo S, Reguenga C, Osorio L, Pereira C, Lopes C, Lima D. DRG11 immunohistochemical expression during embryonic development in the mouse. Developmental Dynamics. 2007;236(9):2653–60.

25. Killackey HP, Fleming K. The role of the principal sensory nucleus in central trigeminal pattern formation. Developmental Brain Research. 1985;22(1):141–5.

26. Dvoryanchikov G, Hernandez D, Roebber JK, Hill DL, Roper SD, Chaudhari N. Transcriptomes and neurotransmitter profiles of classes of gustatory and somatosensory neurons in the geniculate ganglion. Nature Communications. 2017;8.

27. Moore JD, Lindsay NM, Deschenes M, Kleinfeld D. Vibrissa Self-Motion and Touch Are Reliably Encoded along the Same Somatosensory Pathway from Brainstem through Thalamus. Plos Biology. 2015;13(9).

